# The cholesterol pathway of the Golgi stress response induces cell death and transcription of Golgi-related genes through metabolic dysregulation of phosphatidylinositol-4-phosphate

**DOI:** 10.1101/2023.05.18.541279

**Authors:** Kanae Sasaki, Takuya Adachi, Fumi Morishita, Marika Toide, Yuto Watanabe, Hajime Tajima Sakurai, Sadao Wakabayashi, Satoshi Kusumi, Toshiyuki Yamaji, Kaori Sakurai, Daisuke Koga, Kentaro Hanada, Masafumi Yohda, Hiderou Yoshida

## Abstract

The Golgi stress response is an important cytoprotective system that enhances Golgi function in response to cellular demand, while cells damaged by prolonged Golgi stress undergo cell death to ensure the survival of organisms. OSW-1, a natural compound with anticancer activity, acts as a potent inhibitor of OSBP that transports cholesterol and phosphatidylinositol-4-phosphate (PI4P) at contact sites between the endoplasmic reticulum and the Golgi apparatus. Previously, we reported that OSW-1 induces the Golgi stress response, resulting in Golgi stress-induced transcription and cell death. However, the underlying molecular mechanism has been unknown. To reveal the mechanism of a novel pathway of the Golgi stress response regulating transcriptional induction and cell death (the cholesterol pathway), we performed a genome-wide knockout screen and found that transcriptional induction as well as cell death induced by OSW-1 was repressed in HeLa cells deficient in factors involved in the PI4P metabolism, such as PITPNB and PI4KB genes. Our data indicate that OSW-1 induces Golgi stress-dependent transcriptional induction and cell death through dysregulation of the PI4P metabolism in the Golgi apparatus.

## Introduction

Each organelle in eukaryotic cells has its own mechanism of the stress response that maintains homeostasis of the organelle to cope with insufficiency of organelle function. For example, the endoplasmic reticulum (ER), where membrane and secretory proteins are synthesized and properly folded, has the mechanism of the stress response called the ER stress response (also called the unfolded protein response) (Karagöz et al., 2019; Ron et al., 2012; Mori et al., 2015; Kimata et al., 2011; Hetz et al., 2020; Wiseman et al., 2022). The mammalian ER stress response consists of three response pathways: the IRE1, ATF6 and PERK pathways, which upregulate the expression of ER chaperones and ER-associated degradation (ERAD) factors and repress translation upon accumulation of unfolded or misfolded proteins in the ER.

The Golgi apparatus is an organelle that executes post-translational modifications including glycosylation and sorting of membrane proteins and secretory proteins. When the production of these proteins in the ER is increased and a majority of them are transported into the Golgi, the function of the Golgi becomes insufficient (Golgi stress). To augment the function and relieve Golgi stress, cells activate the Golgi stress response to upregulate the expression of Golgi-related proteins such as glycosylation enzymes and transport components at the level of transcription (Sasaki and Yoshida, 2015). Previously, we identified three response pathways controlling the Golgi stress response: the TFE3 pathway (Taniguchi et al., 2015), the proteoglycan pathway (Sasaki et al., 2019) and the mucin pathway (Jamaludin et al., 2019), which augment Golgi general function, proteoglycan-type glycosylation and mucin-type glycosylation, respectively.

OSW-1 is a natural anticancer compound isolated from the bulbs of *Ornithogalum saundersiae*, which is a herbaceous plant belonging to Liliaceae family (Kubo et al., 1992; Mimaki et al., 1997). OSW-1 has been shown to inhibit cell proliferation and induce apoptosis in cancer cells with high selectivity (Mimaki et al., 1997). However, the mechanism by which OSW-1 induces cell death has been controversial. Several research groups have reported that OSW-1 depolarizes mitochondrial membranes and induces mitochondria-dependent apoptosis with a rise in cytoplasmic calcium (Zhan et al., 2021), for example, by inhibition of sodium-calcium exchanger 1 (NCX1) (Garcia-Prieto et al., 2013). Other researchers have reported mitochondria-independent apoptosis by OSW-1 (Iguchi et al., 2019; Zhu et al., 2005).

OSBP and OSBP2 were identified as molecular targets of OSW-1 (Burgett et al., 2011). OSBP and OSBP2 belong to a family of lipid transfer proteins highly conserved in eukaryotes (ORPs) and act as transporters of lipids such as cholesterol and phospholipids at membrane contact sites (MCSs) (Nakatsu et al., 2021). OSBP contains a PH domain, FFAT-motif and OSBP-related domain (ORD). OSBP is localized to MCSs between the ER and the Golgi through its PH domain interacting with phosphatidylinositol-4-phosphate (PI4P) and the small GTPase ARF1 at the Golgi membranes and the FFAT-motif interacting with VAMP-associated protein A/B (VAPA/B) at the ER membranes (Nakatsu et al., 2021). OSBP transfers PI4P from the Golgi to the ER, while it transports cholesterol from the ER to the Golgi. This PI4P consumption at the Golgi membranes enables OSBP to drive the delivery of cholesterol from the ER (Mesmin et al., 2017). OSW-1 inhibits the exchange transport of cholesterol and PI4P between the ER and the Golgi (Albulescu et al., 2017). OSBP2 is the protein most closely related to OSBP and also has a PH domain, FFAT-motif and ORD as well as OSBP. OSBP2 forms a heterodimer with OSBP and is localized to the Golgi (Wyles et al., 2007), though the function of OSBP2 has not been clarified yet in terms of the lipid transfer at MCSs between the ER and the Golgi.

We have previously discovered that OSW-1 activates the TFE3 pathway of the Golgi stress response in HeLa cells (Kimura et al., 2019), while Oh-hashi and colleagues have recently reported that OSW-1 induces atypical Golgi stress and autophagy in Neuro2a cells (Oh-hashi et al., 2023). Therefore, we speculated that the Golgi stress response also contributes to OSW-1-induced transcription and cell death in addition to mitochondrial depolarization. In this report, to clarify the primary cause of cell death induced by OSW-1, we searched for genes essential for the cell death using the Genome-wide CRISPR-Cas9 knockout (GeCKO) system.

## Results

### Inhibition of OSBP and OSBP2 by OSW-1 induces transcription of OSBP2 through the cholesterol pathway of the Golgi stress response

We previously found that the inhibition of OSBP and OSBP2 by OSW-1 increases the transcription of target genes of the TFE3 pathway of the Golgi stress response such as GM130 and GCP60 (Kimura et al., 2019). To get a comprehensive view of the transcriptome of cells treated with OSW-1, we performed RNA sequencing with next-generation DNA sequencing (NGS) technology. Total RNA was prepared from HeLa cells treated with OSW-1 and subjected to RNA sequencing (data not shown) and qRT-PCR experiment (Fig. 1A and 1B). Interestingly, the transcription of OSBP2 was increased upon OSW-1 treatment (Fig. 1B), whereas that of OSBP was not affected (Fig. 1A). Suppression of OSBP expression with siRNA of OSBP (siOSBP) also resulted in the increase of OSBP2 mRNA, indicating that inhibition of OSBP and OSBP2 by OSW-1 induces transcription of OSBP2 (Fig. 1C and 1D). Since OSBP2 is not a target gene of the TFE3 pathway (Oku et al., 2011), another response pathway of the Golgi stress response induces transcription of OSBP2. We named this novel response pathway the cholesterol pathway.

**Figure 1.**
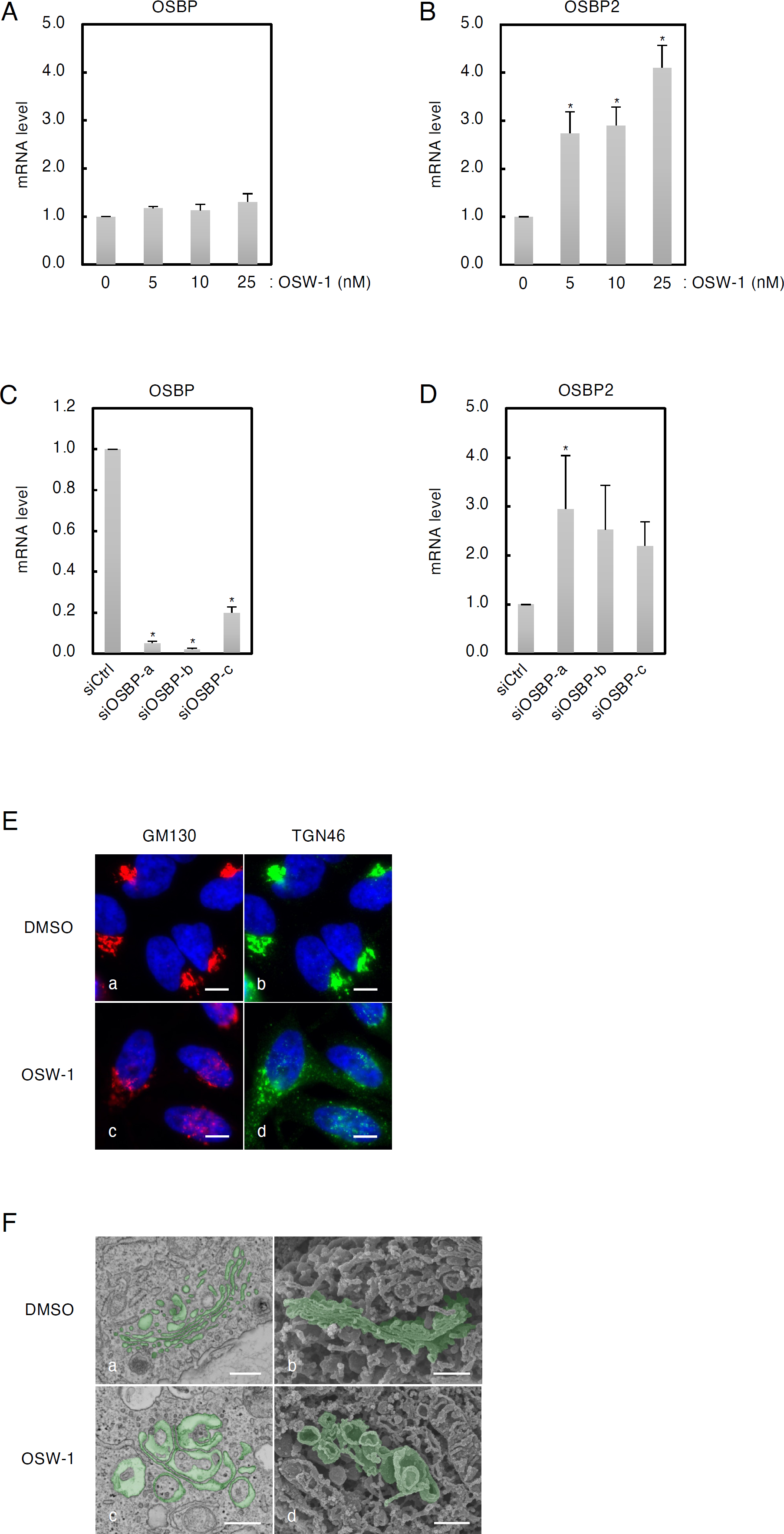
The effect of OSW-1 treatment on transcription and the morphology of the Golgi. (A-D) Quantitative real-time PCR (qRT-PCR) analysis of OSBP and OSBP2 mRNA. RNA extracted from HeLa cells (A and B) or OSBP-knockdown cells (C and D) treated with or without OSW-1 was subjected to qRT-PCR. All mRNA levels were normalized to GAPDH mRNA, which was used as a reference. *, p < 0.05 *versus* each control (analysis of variance followed by Dunnett’s test; mean ± S.E.; n = 3). (E) Immunofluorescence microscopic analysis of the Golgi. Wt HeLa cells were treated with or without 5 nM OSW-1 for 18 h and stained with murine anti-GM130 mAb (red), sheep anti-TGN46 pAb (green), and with DAPI (blue). Scale bars, 10 μm. (F) Electron microscopy analysis of the Golgi. HeLa cells were treated without (a-c) or with 5 nM OSW-1 (d-f) for 18 h, and subjected to SEM analysis. Scale bars, 500 nm.

Previously, it was reported that OSW-1 treatment induces fragmentation of the Golgi (Burgett et al., 2011). As shown in Fig. 1E, we also confirmed Golgi fragmentation by OSW-1 with immunofluorescence analysis using anti-GM130 and anti-TGN46 antibodies in HeLa cells (panels c-d). To precisely analyze microstructural changes of the Golgi by OSW-1, we performed scanning electron microscopy (SEM) of HeLa cells treated with OSW-1 for 18 h (Fig. 1F). Each Golgi cisterna was flat and orderly in HeLa cells without OSW-1 treatment (panels a and b), whereas cells treated with OSW-1 showed swelling and vacuolation of most cisternae (panels c and d). We confirmed that these swollen membrane structures were originated from cisternae of the Golgi using the immunoelectron microscopy method (supplemental Fig. 1). As shown in Fig. 1E (panels a and b) and supplemental movie 1, a set of the Golgi stack was positioned near the nucleus under the normal condition. On the other hand, there were several fragmented Golgi stacks and many vacuoles in OSW-1-treated cells (Fig. 1E panels c, d and supplemental movie 2).

### Screening of genes involved in cell death induced by OSW-1

It was previously reported that OSW-1 induces cell death (Mimaki et al., 1997), and that the Golgi stress response is involved in OSW-1-induced cell death (Kimura et al., 2019; Oh-hashi et al., 2023). However, it has been elusive how the Golgi stress response induces cell death. To identify genes responsible for the OSW-1-induced cell death, we performed GeCKO screening using a GeCKO v2 pooled library A. A total of 125 sgRNAs (corresponding to 64 genes) were enriched in the range of 10- to 681-fold (Fig. 2A), which included five genes related to PI4P metabolism in the ER or the Golgi (Fig. 2B). PITPNB catalyzes the transfer of phosphatidylinositol (PI) from the ER to the Golgi (Cockcroft et al., 2007). CDIPT is CDP-diacylglycerol-inositol 3-phosphatidyl transferase, which catalyzes the *de novo* synthesis of PI from CDP-DAG (CDP-diacylglycerol) (Antonsson, 1994; Lykidis et al., 1997). PI4K2A and PI4KB are PI 4-kinases, which produce PI4P from PI. Both proteins are mainly localized in the *trans* Golgi stacks and TGN (hereafter we use the term “*trans* Golgi regions” to represent the *trans* Golgi stacks and TGN) (Wong et al., 1997; Antonietta et al., 2005). C10orf76 is involved in the activation of PI4KB (McPhail et al., 2020; Mizuike et al., 2023). This result suggests that PI4P metabolism in the *trans* Golgi regions is closely related to OSW-1-induced cell death.

**Figure 2.**
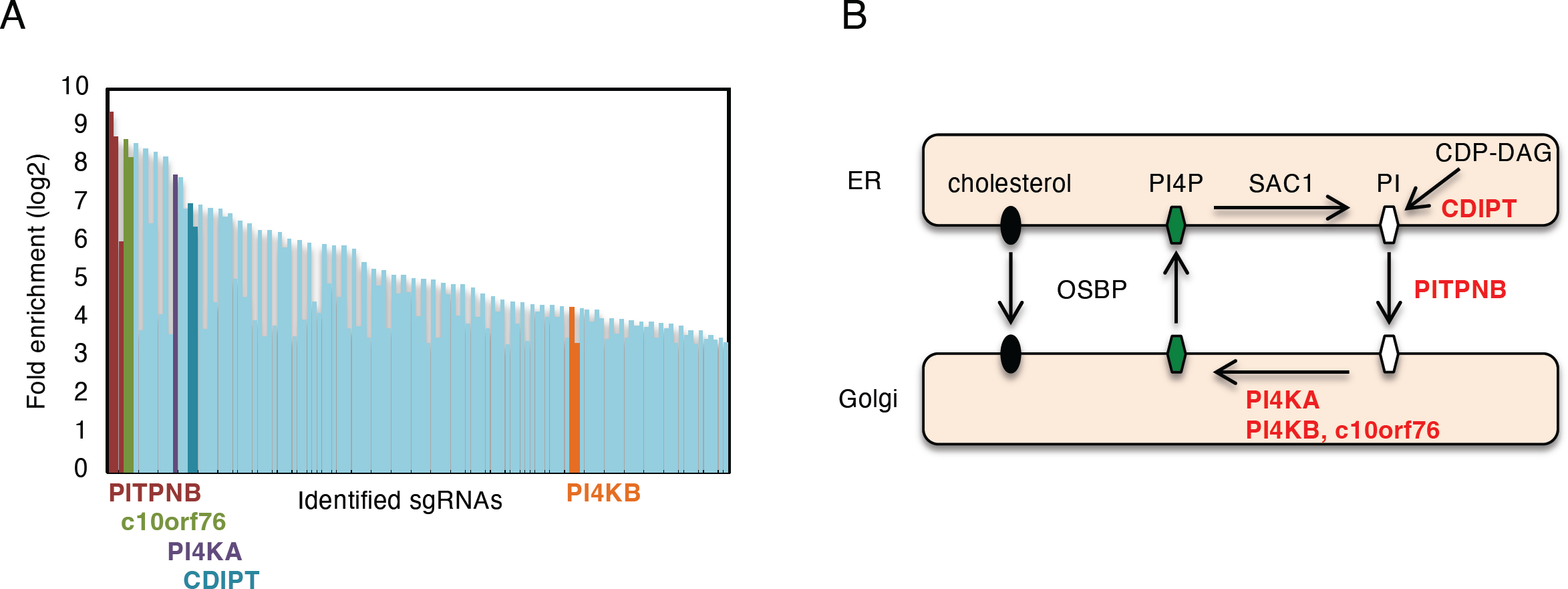
Identification of genes involved in cell death induced by OSW-1 using genome-wide CRISPR-Cas9 knockout screening. (A) Identification of sgRNAs enriched in the OSW-1-resistant HeLa cells. Fold enrichment corresponds to the average of two independent experiments. Red, green, purple, blue and orange bars show 3 sgRNAs targeting the PITPNB gene, 2 sgRNAs targeting to c10orf76 gene, a single sgRNA targeting to PI4K2A gene, 2 sgRNAs targeting the CDIPT gene and 2 sgRNAs targeting the PI4KB gene, respectively. (B) A schematic presentation of PI4P metabolism in the ER and the Golgi. Genes enriched in the screen are shown in red.

### Either PITPNB or PI4KB gene is indispensable for cell death induced by OSW-1

The GeCKO v2 pooled library A contains 3 sgRNAs targeting PITPNB, all of which were highly enriched (Fig. 2A), suggesting that PITPNB is the most promising candidate. We established two stable cell lines of PITPNB-KO cells (PITPNB-KO#2 and #6). To check the expression of PITPNB protein in these clones, we made immunoblotting with anti-PITPNB antibody (Fig. 3A). Although the protein levels of PITPNB in PITPNB-KO#2 and #6 cells were remarkably reduced compared to wild-type (wt) HeLa cells, we detected faint bands at the same position as those of PITPNB proteins in wt cells (lanes 3 and 5). To confirm that the PITPNB gene was completely disrupted in PITPNB-KO#2 and #6 cells, we checked nucleotide sequences of the second exon of the PITPNB gene in these cells, which contains a target sequence of an sgRNA used to establish PITPNB KO cell lines (Fig. 3B). PITPNB-KO#2 showed the deletion of two sequential nucleotides (2 nt) upstream to the sgRNA target region, which caused a frameshift of translation and loss of its mature protein. PITPNB-KO#6 showed two patterns of genome sequences: one was a 1 bp deletion resulting in a frameshift, and the other was 6 bp deletion, which did not cause a frameshift. Although these results could not exclude the possibility that each clone had at least one copy with no editing or no frameshift, cell death by OSW-1 was markedly repressed in PITPNB-KO#2 and #6 compared to wt cells (Fig. 3C). Importantly, the ectopic expression of PITPNB (Fig. 3A, lanes 4 and 6) rescued the sensitivity to OSW-1 in PITPNB-KO cells (Fig. 3D, lanes 8 and 12), indicating that the resistance to OSW-1 is ascribed to the loss of PITPNB, not to any off-targeting effects of gene disruption experiments. To rule out the possibility that the cell death machinery itself is lost in PITPNB-KO#2 and #6 cells, we examined the sensitivity of PITPNB-KO#2 and #6 cells to other stress inducers (Fig. 3E and 3F). Thapsigargin, an inducer of the ER stress, killed PITPNB-KO#2 and #6 cells at the same sensitivity as wt cells, verifying that the cell death machinery itself is intact in PITPNB-KO#2 and #6 cells. Sensitivity to monensin, an activator of the TFE3 pathway of the Golgi stress response, was the same for both wt and PITPNB-KO cells. This suggested that the cholesterol pathway activated by the OSW-1 is distinct from the TFE3 pathway.

**Figure 3.**
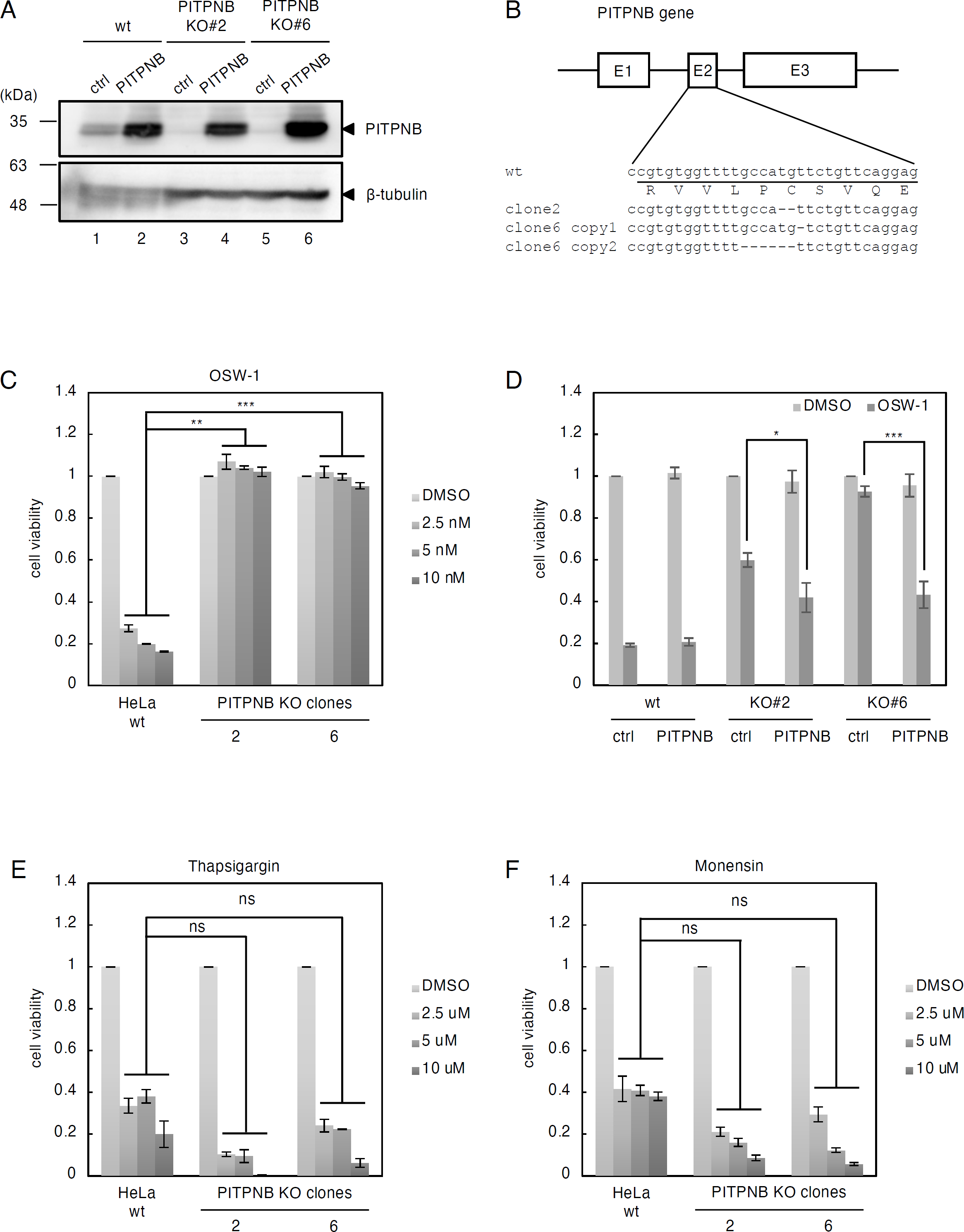
OSW-1-induced cell death in PITPNB knockout cells. (A) Immunoblot analysis of PITPNB protein. Wt HeLa cells and two individual clones (#2 and #6) of PITPNB KO cells were transfected with an empty vector (ctrl) or an expression plasmid of PITPNB (PITPNB). β-tubulin was used as a loading control. (B) Genomic sequences of exon 2 of human PITPNB gene in wt HeLa cells and PITPNB KO clones (#2 and #6). Deletion of two nucleotides was confirmed in both copies of the PITPNB gene in clone 2, resulting in a frameshift. Deletion of one or six nucleotides was confirmed in each copy of the PITPNB gene in clone 6. (C, E and F) Cell viability assay. Wt HeLa cells and PITPNB KO clones (#2 and #6) were treated with or without the indicated concentrations of OSW-1, thapsigargin or monensin for 48 h, and subjected to cell viability assay. **, p < 0.01, ***, p < 0.001 versus wt cells supplemented with the same concentration of each reagent (the Bonferroni-corrected t-test; mean ± S.E.; n = 3). ns, not significant. (D) Rescue experiments of PITPNB KO cells. Wt HeLa cells and PITPNB KO clones (#2 and #6) were transfected with an empty vector (ctrl) or an expression plasmid of PITPNB (PITPNB), treated with or without 5 nM OSW-1 for 48 h, and subjected to cell viability assay. *, p < 0.05, ***, p < 0.001 (the Bonferroni-corrected t-test; mean ± S.E.; n = 3).

Two stable cell lines of PI4KB-KO cells (PI4KB-KO #5 and #7) were also established. Immunoblotting analysis showed that the expression of PI4KB protein was completely lost (Fig. 4A, lanes 2 and 3). Genomic DNA analysis revealed that both alleles of the PI4KB gene were disrupted with one and two nucleotide deletions, which cause frameshift (Fig. 4B). Cell death by OSW-1 was suppressed in PI4KB-KO #5 and #7 cells compared to wt cells (Fig. 4C). The cell death machinery was not disrupted in PI4KB-KO #5 and #7 cells since sensitivity to thapsigargin and monensin is the same for PI4KB-KO #5 and #7 and wt cells (Fig. 4D and 4E). These results lead us to the previously unrecognized possibility that PI4P levels in the *trans* Golgi regions may be responsible for cell death caused by OSW-1.

**Figure 4.**
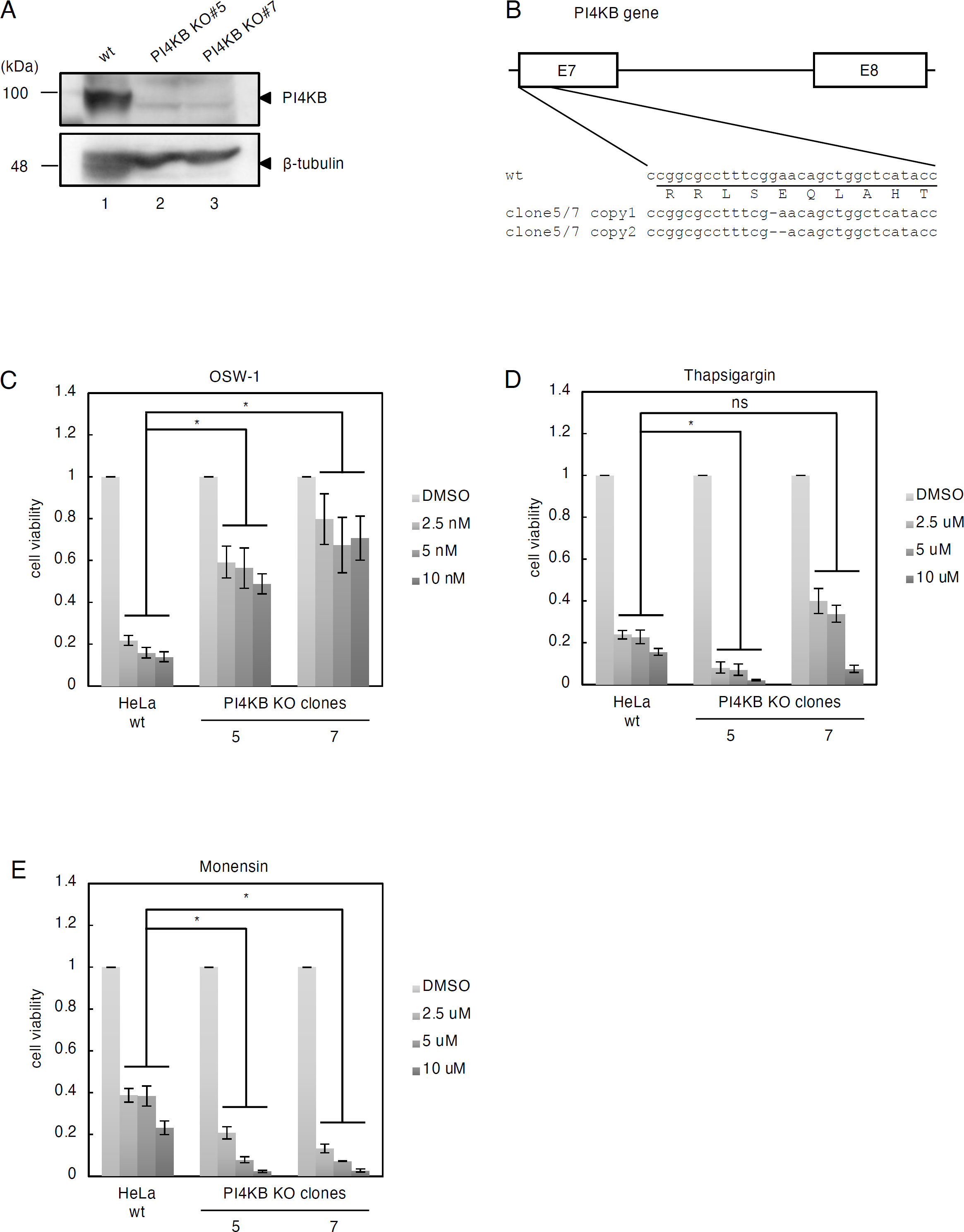
OSW-1-induced cell death in PI4KB knockout cells. (A) Immunoblot analysis of PI4KB protein. Wt HeLa cells and two individual clones (#5 and #7) of PI4KB KO cells were transfected with an empty vector (ctrl) or an expression plasmid of PI4KB (PI4KB). β-tubulin was used as a loading control. (B) Genomic sequences of exon 7 of human PI4KB gene in wt HeLa cells and PI4KB KO clones (#5 and #7). Deletion of one or two nucleotides was confirmed in both copies of the PI4KB gene in clones 5 and 7, resulting in a frameshift. (C-E) Cell viability assay. Wt HeLa cells and PI4KB KO clones (#5 and #7) were treated with or without the indicated concentrations of OSW-1, thapsigargin or monensin for 48 h, and subjected to cell viability assay. *, p < 0.05 versus wt cells supplemented with the same concentration of each reagent (the Bonferroni-corrected t-test; mean ± S.E.; n = 3). ns, not significant.

### Either PITPNB or PI4KB is indispensable for the transcriptional induction of target genes of the cholesterol pathway in response to Golgi stress

RNA sequencing on HeLa cells treated with OSW-1 revealed the transcription of OSBP2 (Fig. 1B and 5B) and FABP3 (Fig. 5B), fatty acid-binding protein 3, was increased upon OSW-1 treatment. To investigate whether PI4P levels in the *trans* Golgi regions are responsible for the transcriptional induction of OSBP2 and FABP3, we examined their transcription in PITPNB- and PI4KB-KO cells by qRT-PCR experiments (Fig. 5). As expected, the transcriptional induction of OSBP2 and FABP3 was diminished in PITPNB-or PI4KB-KO cells (Fig. 5A-D). These data suggest that transcriptional induction of OSBP2 and FABP3 by the cholesterol pathway is triggered depending on PI4P levels in the *trans* Golgi regions as observed in cell death induced by OSW-1.

**Figure 5.**
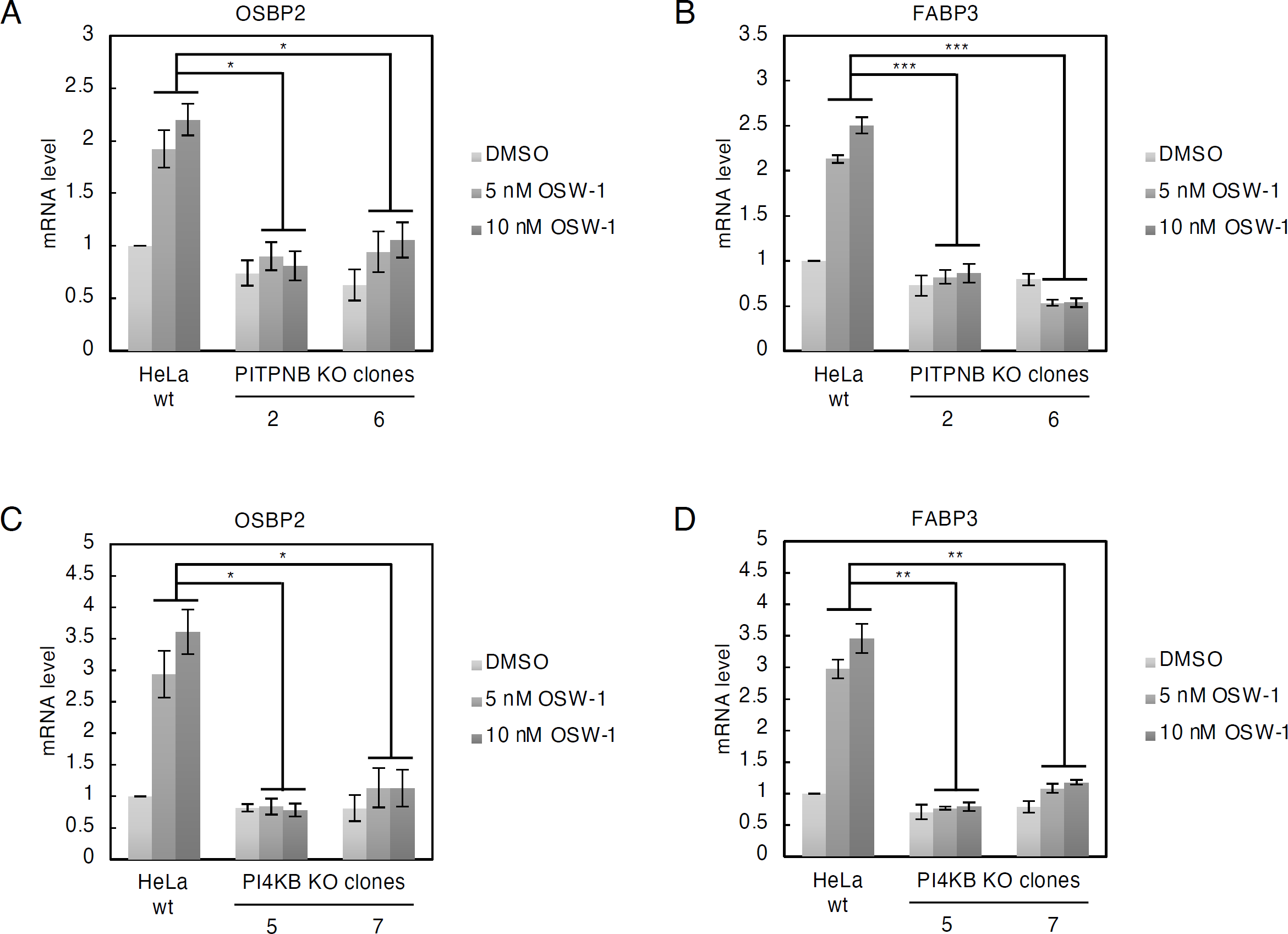
Transcriptional induction of OSBP2 and FABP3 genes by OSW-1 in PITPNB KO and PI4KB KO cells. (A-D) qRT-PCR analysis of OSBP and FABP3 mRNA. HeLa wt cells, PITPNB KO clones (#2 and #6) and PI4KB KO clones (#5 and #7) were treated with or without 5 nM OSW-1 for 18 h, and subjected to qRT-PCR analysis. All mRNA levels were normalized to GAPDH mRNA, which was used as a reference. *, p < 0.05, **, p < 0.01, ***, p < 0.001 (the Bonferroni-corrected t-test; mean ± S.E.; n = 3).

### Either PITPNB or PI4KB is indispensable for Golgi fragmentation induced by OSW-1

Previously, it was reported that OSW-1 treatment induces Golgi fragmentation before cell death. To further investigate whether PI4P levels in the *trans* Golgi regions are responsible for Golgi fragmentation caused by OSW-1, we performed immunostaining using antibodies against GM130 and TGN46, which are a *cis*-Golgi marker and a TGN marker, respectively (Fig. 6). In wt HeLa cells, GM130 and TGN46 were accumulated in the perinuclear region under the normal condition (Fig. 6A, panels a-1 and a-6), whereas they began to disperse about 6 h later upon the treatment of OSW-1 (panels a-3∼5 and a-8∼10). Unexpectedly, we did not observe the remarkable dispersion of GM130 and TGN46 even 18 h after OSW-1 treatment in PITPNB or PI4KB KO cell lines (panels b-3∼5, b-8∼10, c-3∼5, c-8∼10, d-3∼5, d-8∼10, e-3∼5 and e-8∼10). Statistically significant decreases in the percentage of cells showing *cis*-Golgi fragmentation by OSW-1 were observed in PITPNB or PI4KB KO cells compared to wt cells (ratio of *cis*-Golgi-fragmented cells: wt, 86 ± 3.3%; PITPNB KO#2, 6.8 ± 3.2%; PITPNB KO#6, 25 ± 9.7%; PI4KB KO#5, 16 ± 3.9%; PI4KB KO#7, 5.2 ± 3.2%) (Fig. 6B). Similarly, decreases in the percentage of cells showing TGN fragmentation by OSW-1 were observed in PITPNB or PI4KB KO cells (ratio of TGN-fragmented cells: wt, 99 ± 0.56%; PITPNB KO#2, 39 ± 5.9%; PITPNB KO#6, 30 ± 2.5%; PI4KB KO#5, 55 ± 16%; PI4KB KO#7, 26 ± 15%) (Fig. 6C). These data indicate that both PITPNB and PI4KB are indispensable for Golgi fragmentation induced by OSW-1.

**Figure 6.**
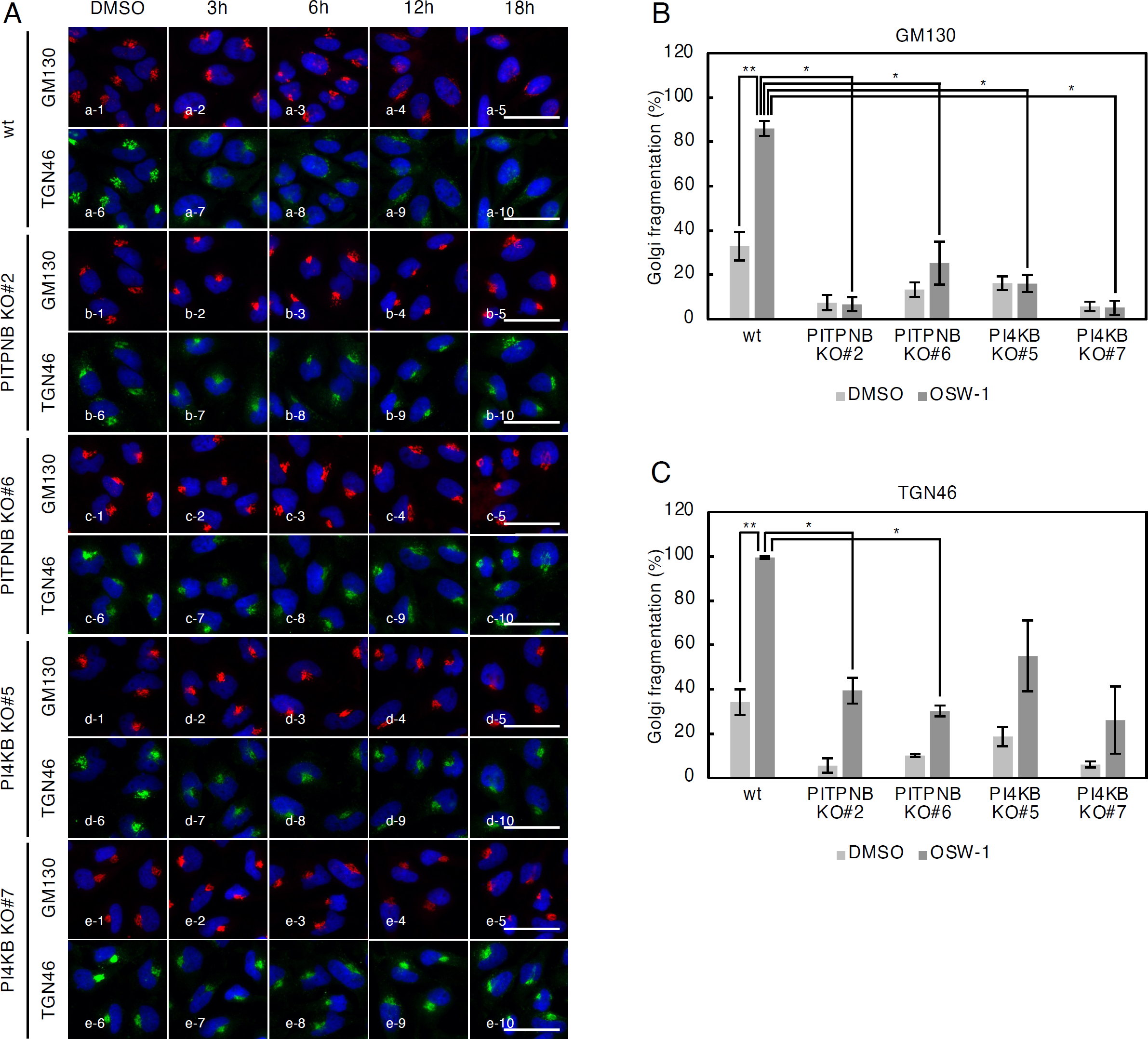
Golgi fragmentation by OSW-1 in PITPNB or PI4KB KO cells. (A) Immunofluorescence microscopic analysis of HeLa wt cells, PITPNB KO clones (#2 and #6) and PI4KB KO clones (#5 and #7) treated with or without 5nM OSW-1 for the indicated time periods. Cells were stained with murine anti-GM130 mAb (red), sheep anti-TGN46 pAb (green), and DAPI (blue). Scale bars, 50 μm. (B and C) Quantification of Golgi fragmentation in wt HeLa cells, PITPNB KO clones (#2 and #6) and PI4KB KO clones (#5 and #7) treated with or without 5 nM OSW-1 for 18 h. The percentage of cells with fragmented Golgi was calculated using images immunostained with anti-GM130 mAb or anti-TGN46 pAb. *, p < 0.05, **, p < 0.01 (the Bonferroni-corrected t-test; mean ± S.E.; n = 3).

### OSW-1 abolishes glycosylation in the Golgi depending on PI4P levels in the *trans* **Golgi regions**

To investigate the events in the Golgi fragmented by OSW-1 treatment, immunoblotting analysis was performed using anti-GM130 and TGN46 antibodies. The molecular weight of GM130 did not change after OSW-1 treatment in wt HeLa cells (Fig. 7A and 7B, upper panel, lanes 1-4), whereas that of TGN46 markedly shifted from 100 kDa to 75 kDa (middle panel, lanes 1-4).

**Figure 7.**
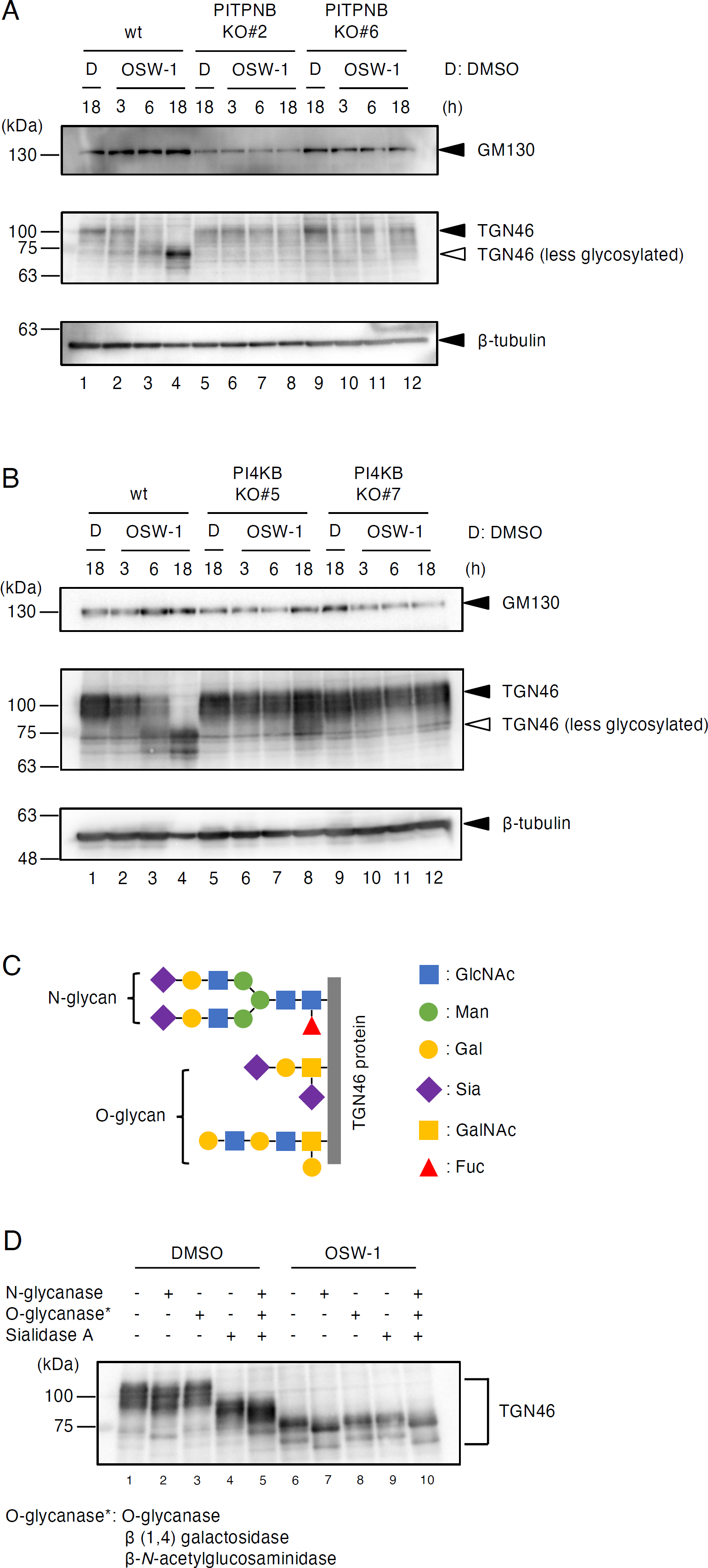
Glycosylation in the Golgi of PITPNB and PI4KB KO cells. (A and B) Immunoblot analysis of GM130 and TGN46 proteins. Whole cell lysates prepared from wild type and PITPNB KO cells (A) or PI4KB KO cells (B), treated with or without 5 nM OSW-1 for 18 h, were subjected to immunoblot analysis. β-tubulin was used as a loading control. (C) A schematic presentation of the glycosylation pattern on TGN46. *N*- and *O*-linked glycans including sialic acids are attached to TGN46 (Durin et al., 2022; van Galen et al., 2014). (D) Deglycosylation assay of the TGN46 protein in wt HeLa cells treated with or without 5 nM OSW-1 for the indicated time periods. Deglycosylation enzymes used in this assay were as follows: N-glycanease, O-glycanase mix (O-glycanase, β (1-4) galactosidase and β -N-acetylglucosaminidase), and sialidase A.

To investigate whether the molecular weight shift of TGN46 caused by OSW-1 treatment was due to insufficient glycosylation, we examined the glycosylation state of TGN46, which has been reported to be a glycoprotein with *N*- and *O*-linked glycans (Durin et al., 2022; van Galen et al., 2014). Whole cell lysates of wt HeLa cells were treated with *N*-glycanase to remove *N*-glycan, *O*-glycanase, β (1,4) galactosidase and β-*N*-acetylglucosaminidase to degrade *O*-glycan, and sialidase A to remove sialic acid, and subjected to immunoblotting (Fig. 7C). The treatment of these deglycosylation enzymes reduced the molecular weight of TGN46 (lanes 1 and 5), indicating that TGN46 is highly glycosylated. Even after OSW-1 treatment, the molecular weight of TGN46 was further reduced by *N*-glycanase treatment (lanes 6 and 7), indicating that the *N*-glycosylation, which occurs in the ER, was not affected by OSW-1. In contrast, the molecular weight of TGN46 was not changed by *O*-glycanase, β (1,4) galactosidase, β-*N*-acetylglucosaminidase and sialidase A treatments (lanes 6, 8 and 9). This indicates that the sialidation and *O*-linked glycosylation, which occurs in the Golgi, was inhibited by OSW-1. Interestingly, the molecular weight of TGN46 returned to normal in PITPNB-KO and PI4KB-KO cells (Fig. 7A and 7B, lanes 5-12), suggesting that OSW-1 represses Golgi functions such as glycosylation depending on PI4P levels in the *trans* Golgi regions.

## Discussion

In this study, we revealed that OSW-1 treatment evokes fragmentation of the Golgi, inhibition of glycosylation in the Golgi, transcriptional induction of OSBP2 and cell death. Disruption of genes involved in PI4P synthesis in the *trans* Golgi regions including PI4KB and PITPNB canceled these effects of OSW-1. Based on these observations, we propose that OSW-1 induces transcription and cell death by activating a novel response pathway of the Golgi stress response, the cholesterol pathway, through dysregulation of the PI4P metabolism in the *trans* Golgi regions (Fig. 8).

**Figure 8.**
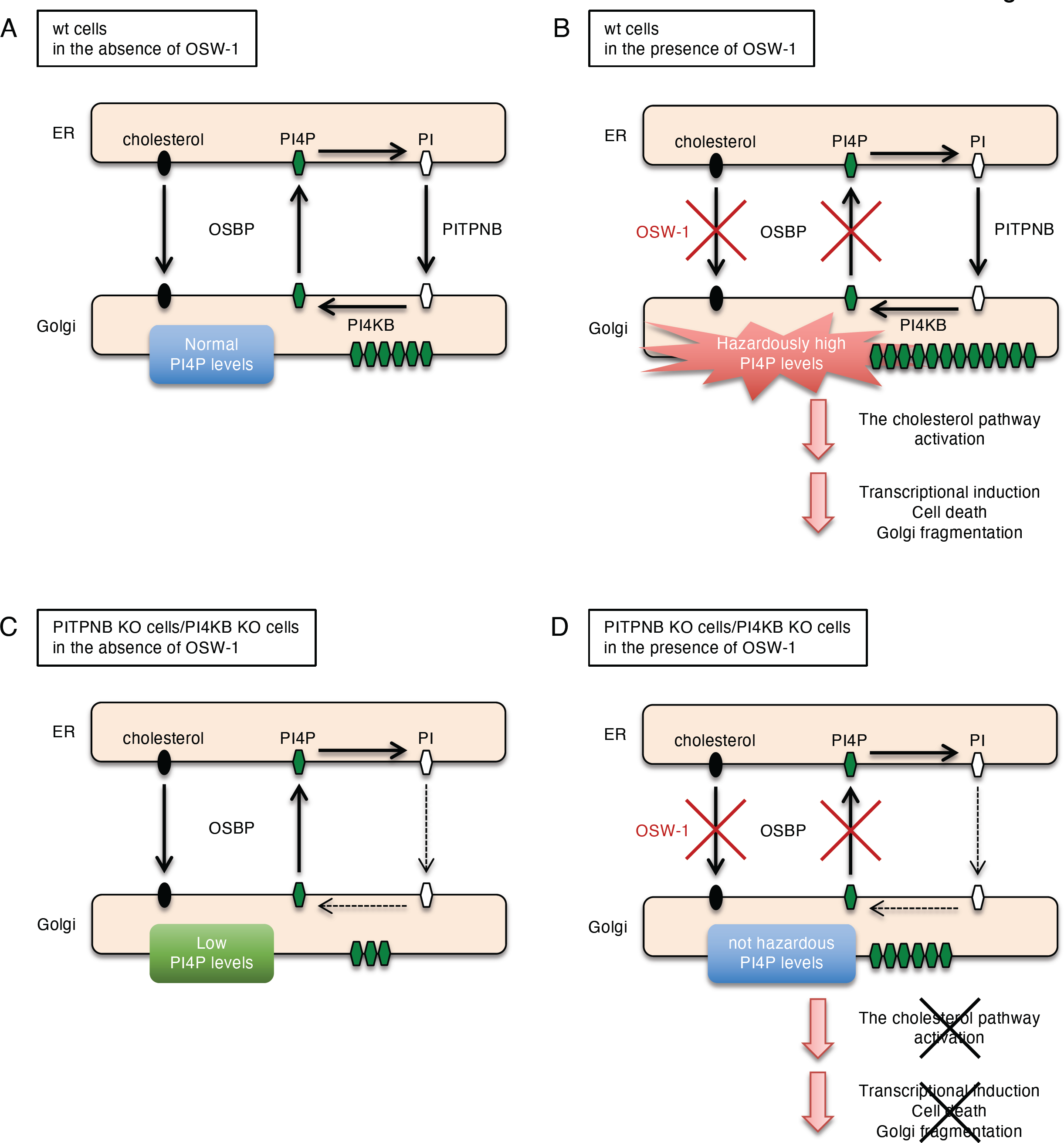
Our working hypothesis for the activation of the cholesterol pathway of the Golgi stress response. In wild-type cells, treatment of OSW-1 inhibits OSBP, leading to the accumulation of PI4P in the *trans* Golgi regions, which causes transcriptional activation, cell death and Golgi fragmentation through the activation of the cholesterol pathway. In contrast, PITPNB or PI4KB KO cells are presumed to exhibit lower levels of PI4P upon treatment of OSW-1 since the PITPNB and PI4KB genes are involved in PI4P synthesis. Therefore, in PITPNB or PI4KB KO cells, OSW-1 treatment seems not to induce accumulation of PI4P at concentrations high enough to activate the cholesterol pathway. As a result, cell death is repressed in these KO cells.

We performed RNA sequencing (data not shown) and qRT-PCR analysis (Fig. 1A-D) of HeLa cells treated with or without OSW-1, and found that transcription of the OSBP2 and FABP3 genes is increased upon OSW-1 treatment (Fig. 5A and 5B), while we did not detect an increase of OSBP mRNA expression (Fig. 1A). The PH and ORD domains of OSBP2 bind to PI4P and cholesterol, respectively (Charman et al., 2014), and OSBP2 is localized to the Golgi by forming a heterodimer with OSBP (Wyles et al., 2007), suggesting that OSBP2 transports cholesterol and PI4P between the ER and the Golgi like OSBP, though it should be determined by direct experiments. Since OSW-1 binds to the ORD of OSBP with higher affinity than that of OSBP2 (Roberts et al., 2019), OSBP2 may not be completely inhibited by OSW-1. Thus, it is possible that OSBP is a constitutively expressed type, while OSBP2 is an inducible one to compensate for the cholesterol/PI4P transfer activity between the ER and the Golgi. FABP3 belongs to the FABP family, which consists of nine members that are expressed in a tissue-specific manner, and is highly expressed in the heart and skeletal muscles (Furuhashi et al., 2008). FABPs bind to long-chain fatty acids and are considered to play a role in the transport of lipids to intracellular compartments such as lipid droplets and organelles for storage, metabolism and signaling, and extracellular regions for signaling (Furuhashi et al., 2008). FABP3 is upregulated especially in aged skeletal muscles, reduces membrane fluidity by increasing lipid saturation and induces ER stress (Lee et al., 2020). The upregulation of FABP3 expression observed here may reflect a disturbance of lipid composition in the Golgi membrane. The mammalian Golgi stress response consists of several response pathways including the TFE3 pathway, the proteoglycan pathway, the mucin pathway, the CREB3 pathway and the HSP47 pathway (Sasaki and Yoshida, 2015). Since OSBP2 and FABP3 are not included in the list of target genes of these well-known pathways, the transcriptional induction of the OSBP2 and FABP3 is thought to be regulated by a novel response pathway of the Golgi stress response, that is, the cholesterol pathway, which is activated by metabolic dysregulation of PI4P in the *trans* Golgi regions.

PITPNB, PI4KB, PI4K2A, CDIPT and C10orf76 were identified as candidates essential for cell death induced by OSW-1. All of these proteins are involved in the synthesis of PI4P in the in the *trans* Golgi regions. PITPNB encodes PITPβ, which belongs to the PITP protein family with one common PITP domain and has exchange activity of phosphatidylinositol (PI) / phosphatidylcholine (PC) or PI/phosphatidic acid (PA) between different organelles (Cockcroft et al., 2020). The PITP family includes five proteins: PITPα, PITPβ, PITPNM1, PITPNM2 and PITPNC1. Among them, PITPα and PITPβ share the most homologous common structure and are ubiquitously expressed in mammals. Xie et al. found that both PITPα and PITPβ regulate the production of the PI4P pool at the Golgi, which is essential for the normal development of the neocortex in mammalian embryos (Xie et al., 2018). However, our GeCKO screen did not identify any genes encoding members of the PITP protein family other than PITPβ. It may be because PITPβ shuttles between the ER and the Golgi, while PITPα resides in the cytosol (Shadan et al., 2008). Although PITPNC1 has also been shown to maintain a pool of PI4P in the Golgi, there are no reports comparing the contribution of these proteins to the production of PI4P. PITPβ, together with regulators such as OSBP, PI4KB, and Sac1, is localized to the MCS between the ER and the Golgi and is presumed to actively contribute to the regulation of PI4P levels in the Golgi of mammalian cells.

PI4KB and PI4K2A are PI 4-kinases that generate a pool of PI4P in the Golgi (Wong et al., 1997; Wang et al., 2003). CDIPT protein catalyzes the *de novo* synthesis of PI (Antonsson, 1994; Lykidis et al., 1997). On the other hand, C10orf76 forms a direct complex with PI4KB and contributes PI4KB activation via the recruitment of ARF1-GTP to the Golgi. KO of C10orf76 reduces PI4P levels in the Golgi in HAP1 cells (Blomen et al., 2015; McPhail et al., 2020) and in HeLa cells (Mizuike et al., 2023). From these, it is inferred that inhibition of PI4P transport from the *trans* Golgi regions to the ER by OSW-1 leads to excessive accumulation of PI4P in the *trans* Golgi regions and drastic change in the lipid composition of the Golgi membrane, resulting in the collapse of the Golgi, activation of the Golgi stress response, and finally induction of OSBP2 transcription and cell death in wt cells (Fig. 8B). On the other hand, PI4P levels in PI4KB, PI4K2A, PITPNB, CDIPT or C10orf76 KO cells treated with OSW-1 should be lower than those in wt cells and could not activate the cholesterol pathway of the Golgi stress response (Fig. 8D). Although it has been already reported that a decrease in PI4P levels inhibits Golgi function (Santiago-Tirado and Bretscher, 2011), this is the first report that excessive accumulation of PI4P in the *trans* Golgi regions represses Golgi function.

Although OSW-1 exerts potent and selective anticancer effects, the question of how inhibition of OSBP by OSW-1 leads to cell death specific to cancer cells has remained unresolved for a long time. Increased expression of PI-related factors localized to the Golgi, such as PI4KB, SAC1, GOLPH3 and PITPNC1, is one of the common features of several tumors (Waugh, 2019). For instance, in some tumors, PI4KB is overexpressed due to the multiplication of the PI4KB gene, resulting in the overproduction of PI4P in the *trans* Golgi regions. If PI4P is increased in the *trans* Golgi regions, PI4P-binding proteins like GOLPH3 and PITPNC1 are over-recruited, leading to malignant secretion that can develop angiogenesis, tumor invasion and metastasis. Thus, we speculate that the anticancer effect of OSW-1 is derived from the increase of PI4P levels in the Golgi of cancer cells. This is consistent with our finding that depletion of factors related to PI4P synthesis in the Golgi, such as PITPNB and PI4KB, leads to resistance to OSW-1. In addition to conventional anticancer drugs, the development of cancer medicines with various molecular targets is in progress worldwide. Since cancer cells continuously mutate, change their properties, and eventually acquire drug resistance, further exploration of a wide range of molecular targets to kill cancer cells is essential to solving the problem of drug resistance. In this study, we clearly revealed that OSW-1 is a unique anticancer reagent that targets the Golgi function. Our discovery will not only elucidate the role of the Golgi in the malignant transformation of cancer but it will also lead to the development of Golgi-based cancer therapies in the future.

## Materials and methods

### Antibodies and reagents

The following antibodies were purchased: rabbit anti-PITPNB polyclonal antibody (pAb) and sheep anti-TGN46 pAb from Sigma-Aldrich (St. Louis, MO); rabbit anti-β-tubulin pAb from cell signaling (Danvers, MA); rabbit anti-PI4KB pAb from Abcam (Cambridge, UK) and mouse anti-GM130 monoclonal antibody (mAb) from BD Biosciences (Franklin Lakes, NJ). OSW-1 was synthesized as described previously (Sakurai et al., 2010). Thapsigargin and monensin were obtained from Calbiochem.

### Plasmid construction

Human PITPNB cDNA (OHu06208C) was purchased from GenScript (Piscataway, NJ). To construct plasmids for knockout of PITPNB and PI4KB genes, each oligo DNA fragment for expression of corresponding sgRNA was inserted into the BbsI site of pSpCas9 (BB)-2A-Puro (PX459) V2.0, which was purchased from Addgene (Watertown, MA) (plasmid #62988).

### Cell culture and transfection

HeLa cells were cultured in Dulbecco’s modified Eagle’s medium (glucose at 4.5 g/liter) (Nacalai Tesque, Kyoto, Japan) supplemented with 10% fetal bovine serum (FBS) and Penicillin-Streptomycin Solution (Wako, Osaka, Japan) at 37°C under humidified air containing 5% CO2. Cells were seeded and cultured for 1 day, and then they were transfected with the expression plasmid DNAs by using FuGENE 6 (Promega, Madison, WI).

### CRISPR screen

Two sgRNA-expressing HeLa cell libraries (A-1 and A-2) using the GeCKO v2.0 library (Libraries A) were previously prepared (Yamaji et al., 2019). sgRNA-expressing cells from each cell library were treated with 5 nM OSW-1 for 48 h. Then, cells were washed with DMEM containing 10% FBS and cultured in conventional conditions for 11 days. Surviving cells were re-plated and treated with 5 nM OSW-1 again for 48h. Five days after treatment, cells were trypsinized and frozen as cell pellets. For untreated controls, sgRNA-expressing cells in each cell library were cultured for the same period as OSW-1-treated cells with several passages.

### Isolation of genomic DNA and genome-integrated sgRNA sequencing

Analysis of genome-integrated sgRNAs was performed as described previously (Yamaji et al., 2019). Genomic DNA from frozen cells was purified using the conventional phenol-chloroform method. The genome-integrated sgRNA sequences was amplified using KOD -Plus-Neo (Toyobo, Osaka, Japan) and 100 μg genomic DNA from untreated cells or OSW-1-treated cells as a PCR template. Primer pairs used for the PCR were as follows: TCTTGTGGAAAGGACGAAACACCG and GCCACTTTTTCAAGTTGATAACGGACTAG. The amplicons were resolved by agarose electrophoresis and extracted from the gel using NucleoSpin Gel and PCR Clean-up XS (Macherey-Nagel, Düren, Germany). The extracted DNA was then processed into libraries for sequencing using Ion Plus Fragment Library Kit and Ion Xpress Barcode Adapters 1-16 (Thermo Fisher Scientific, Waltham, MA). Sequencing was performed using an Ion Proton System (Thermo Fisher Scientific), and then data processing and analysis were carried out using a SgRNA Screening v.1.1 custom plugin (Thermo Fisher Scientific).

### Generation of knockout cell lines

Generation of PITPNB and PI4KB knock-out HeLa cells was carried out using the CRISPR/Cas9 system as described previously (Ran et al., 2013). Briefly, HeLa cells were transfected with the plasmid (pSpCas9 (BB)-2A-Puro (PX459) V2.0/PITPNB sgRNA or pSpCas9 (BB)-2A-Puro (PX459) V2.0/PI4KB sgRNA) using FuGENE 6. After 24 h, cells were selected with 1 μg/ml puromycin for 3 days to remove untransfected cells. Subsequently, cells were plated on 96-well plates in media without puromycin, and single colonies were selected.

### Indel analysis

Isolation of genomic DNA for PCR was performed as described previously (Ramlee et al., 2015). PCR was performed with KOD -Plus-Neo (Toyobo), and blunt-end PCR products were treated with 10 x A-attachment (Toyobo) mix to acquire overhanging dA at the 3’-ends. Products with 3’-dA overhangs were cloned into T-Vector pMD19 (Simple) (Takara Bio Inc., Shiga, Japan) to use as a template for sequencing.

### Cell viability assay

Cell viability assay was performed using a Cell Counting Kit-8 (DOJINDO, Kumamoto, Japan) according to the manufacturer’s instructions. Briefly, HeLa cells were treated with 5 nM OSW-1, 1 μM thapsigargin or 10 μM monensin for 48 h and incubated with CCK-8 solution in the culture medium for 4 h at 37°C in a CO2 incubator. The absorbance at 415 nm was measured using a microplate reader, Sunrise (TECAN, Boston, MA).

### Western blot analysis

Western blot analysis was performed as described previously (Yoshida et al., 2009). Whole cell lysates were resolved by SDS-PAGE, transferred to polyvinylidene difluoride (PVDF) membranes (Immobilon-P, Merck Millipore, Burlington, MA), and probed with specific antibodies. Chemiluminescent signals were detected by a WSE-6300 LuminoGraph III (ATTO, Tokyo, Japan) using EZ-ECL chemiluminescence detection kit for HRP (Sartorius, Göttingen, Germany).

### RNA sample preparation and quantitative RT-PCR

Cells were harvested 18h after the treatment with 5 nM OSW-1, and then total RNA was prepared with Sepasol Super G (Nacalai Tesque). RT-PCR and quantitative PCR (qPCR) were performed using an ABI7500 qPCR instrument (Life Technologies, Carlsbad, CA) and a PrimeScript RTreagent kit with gDNA Eraser and SYBR Premix Ex Taq II (TliRNase H Plus) (TaKaRa) and an ABI7500 qPCR instrument (Life Technologies). Primer pairs used for qRT-PCR were as follows: *OSBP2* (GTACCAGACCCTGTCAGCCAAG and GTCAGGGCCAGCTCTGAGAA), *FABP3* (TGCGGGAGCTAATTGATGGAA and TTCTCATAAGTGCGAGTGCAAACTG) and *GAPDH* (TGCACCACCAACTGCTTAGC and GGCATGGACTGTGGTCATGAG).

### Immunofluorescence microscopic analysis

HeLa cells were seeded onto coverslips in 24-well plates, washed with PBS, fixed in 4% (w/v) paraformaldehyde in PBS at room temperature for 15 min, and then permeabilized in 0.2% Triton X-100 in PBS at room temperature for 10 min. After blocking with 10% FBS in PBS at room temperature for 1h, the cells on the coverslips were incubated with primary antibodies at 4°C overnight and then with secondary antibodies (Alexa Fluor 488-labeled donkey anti-sheep IgG and Alexa Fluor 488-labeled donkey anti-sheep IgG) at room temperature for 1 h. Finally, they were mounted with SlowFade Gold antifade reagent (Invitrogen, Carlsbad, CA) and their images were acquired using an Eclipse Ni microscope (Nikon, Tokyo, Japan) and ORCA-ER digital CCD camera (Hamamatsu Photonics, Hamamatsu, Japan). Staining of PI4P was performed as described previously (Hammond et al., 2009).

Quantification of Golgi fragmentation was performed as described previously (Carvou et al., 2010). In brief, the random fields of GM130 or TGN46 images in immunostaining were analyzed using ImageJ Fiji. For each field, GM130 or TGN46 signals were automatically selected by thresholding and normal/unfragmented Golgi (Size: 500-Infinity, Circularity: 0.20-1.00) was counted using the Analyze particles tool.

### Electron microscopic analysis

HeLa cells were treated with or without 5 nM OSW-1 for 18 h and washed with PBS, fixed in 0.5% (w/v) paraformaldehyde and 0.5% glutaraldehyde in PBS at room temperature for 20 min. Cells were embedded in agarose as previously described (Koga et al., 2012. Specimens for the osmium maceration method were prepared as previously described (Koga et al., 2021). Finally, cells were observed in field emission scanning electron microscopes (S4100 and Regulus, Hitachi-High-Tech, Tokyo, Japan). Samples for the serial section SEM were prepared according to our paper (Koga et al., 2021). The sections were observed in an ultra-high-resolution scanning electron microscope (SU-70, Hitachi High-Tech) to acquire sequential images of target cells. After aligning the serial images using computer software (Amira, Thermo Fisher Scientific), target structures such as the Golgi apparatus, nuclei, and cytoplasm were segmented and reconstructed in three dimensions. Immunoelectron microscopy analysis was performed as described previously (Kusumi et al., 2018).

### Deglycosylation assay

Deglycosylation assay was performed as described previously (Durin et al., 2022). In brief, HeLa cells were treated with or without 5 nM OSW-1 for 18 h and lysed in RIPA buffer supplemented with protease inhibitor cocktail, cOmplete, EDTA-free (Roche) and proteins were quantified using BCA Protein Assay Kit (Thermo Fisher Scientific). The reaction of deglycosylation was performed with the Agilent Enzymatic Deglycosylation Kit for N-Linked and Simple O-Linked Glycans (GK80110) and Agilent Extender Kit for Complex O-Linked Glycans (GK80115), according to manufacturer’s instructions (Agilent, Santa Clara, CA).

### Statistical analysis

Statistically significant differences in Fig. 2D were evaluated by two-tailed unpaired Student’s t-test. Those in Fig. 2C, 3C, 4A-H, 5B and 5C were evaluated by two-tailed unpaired Student’s t-test with Bonferroni correction. p<0.05 is considered to be statistically significant.

## Acknowledgements

We thank the members of JSPS Grant “Organelle Zones” for their critical discussions, and Ms. Mikiko Ochiai and Ms. Miyu Sakamoto for their secretarial and technical assistance. This work was supported by JSPS KAKENHI (Grant numbers JP17K15122, JP17J00067, JP21H02435 and JP22K06208, a Grant-in-Aid for Scientific Research on Innovative Areas of MEXT (JP17H06414 and 17H06417) “Organelle Zones”, The Takeda Science Foundation, The Foundation of Kinoshita Memorial Enterprise, AMED-CREST “Proteostasis” and Nanken-Kyoten (Grant No. 2022-Domestic 08), TMDU.

---

**Figure S1.**
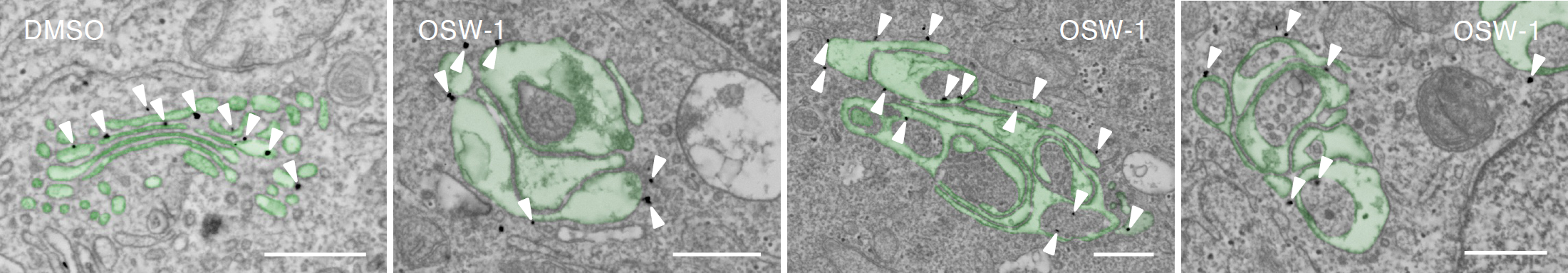
Immunoelectron microscopy analysis of the Golgi. HeLa cells were treated with or without 5 nM OSW-1 for 18 h, stained with murine anti-GM130 mAb, and subjected to SEM observation. *Solid arrowheads*, gold particles of GM130. Scale bars, 500 nm.

**Supplemental movie 1 and 2. 3D reconstruction analysis of the Golgi.**

HeLa cells were treated with or without 5 nM OSW-1 for 18 h, and subjected to 3D reconstruction using serial section SEM images. The Golgi apparatus, nucleus, vacuoles and cytoplasm are shown in green, blue, magenta and orange, respectively.

## Notes

### Competing Interest Statement

The authors have declared no competing interest.

